# Endogenous virus sRNA regulates gene expression following genome shock in tomato hybrids

**DOI:** 10.1101/2021.09.20.461014

**Authors:** Sara Lopez-Gomollon, Sebastian Y. Müller, David C. Baulcombe

## Abstract

Hybridization and environmental stress trigger genome shock that perturbs patterns of gene expression leading to phenotypic changes. In extreme examples it is associated to transposon mobilization and genome rearrangement. Here we discover a novel alternative mechanism in interspecific *Solanum* hybrids in which changes to gene expression were associated with DCL2-mediated small (s)RNAs derived from endogenous (para)retroviruses (EPRVs). Correspondingly, the altered patterns of gene expression overlapped with the effects of *dcl2* mutation and the changes to sRNA profiles involved 22nt species produced in the DCL2 biogenesis pathway. These findings implicate hybridization-induced genome shock leading to EPRV activation and sRNA silencing as causing changes in gene expression. Such hybridization-induced variation in gene expression could increase the range of traits available for selection in natural evolution or in breeding for agriculture.

---

Hybridization within or between species has featured frequently in the evolutionary history of plants and animals. It can have either negative or positive effects on fitness due to genome shock that McClintock associated with transposon activation^1^. In *Drosophila*, for example, P element activation^2,3^ leads to hybrid dysgenesis: a negative effect. The positive effects of genome shock are less well understood and, in principle, they could involve protein-coding genes, regulatory RNA, transposable elements or epigenetic changes^4,5^ leading to changes in gene expression.

To investigate hybrid-induced effects on sRNA and gene expression we crossed *S. lycopersicum* (M82) x *S. pennellii* (LA716) (*lyc* x *pen*) (Fig.1a). The fertile F1 – F4 progeny had variable phenotypes (ExtDataFig.1) that were mostly within the parental ranges and transient in the F1 but some were transgressive (beyond the parental range) and maintained into the F4 (ExtDataFig.1b). This level of phenotypic variation is typical of hybrid populations and justified the use of these hybrids for a more detailed analysis of gene expression and sRNAs.

**Figure 1.**
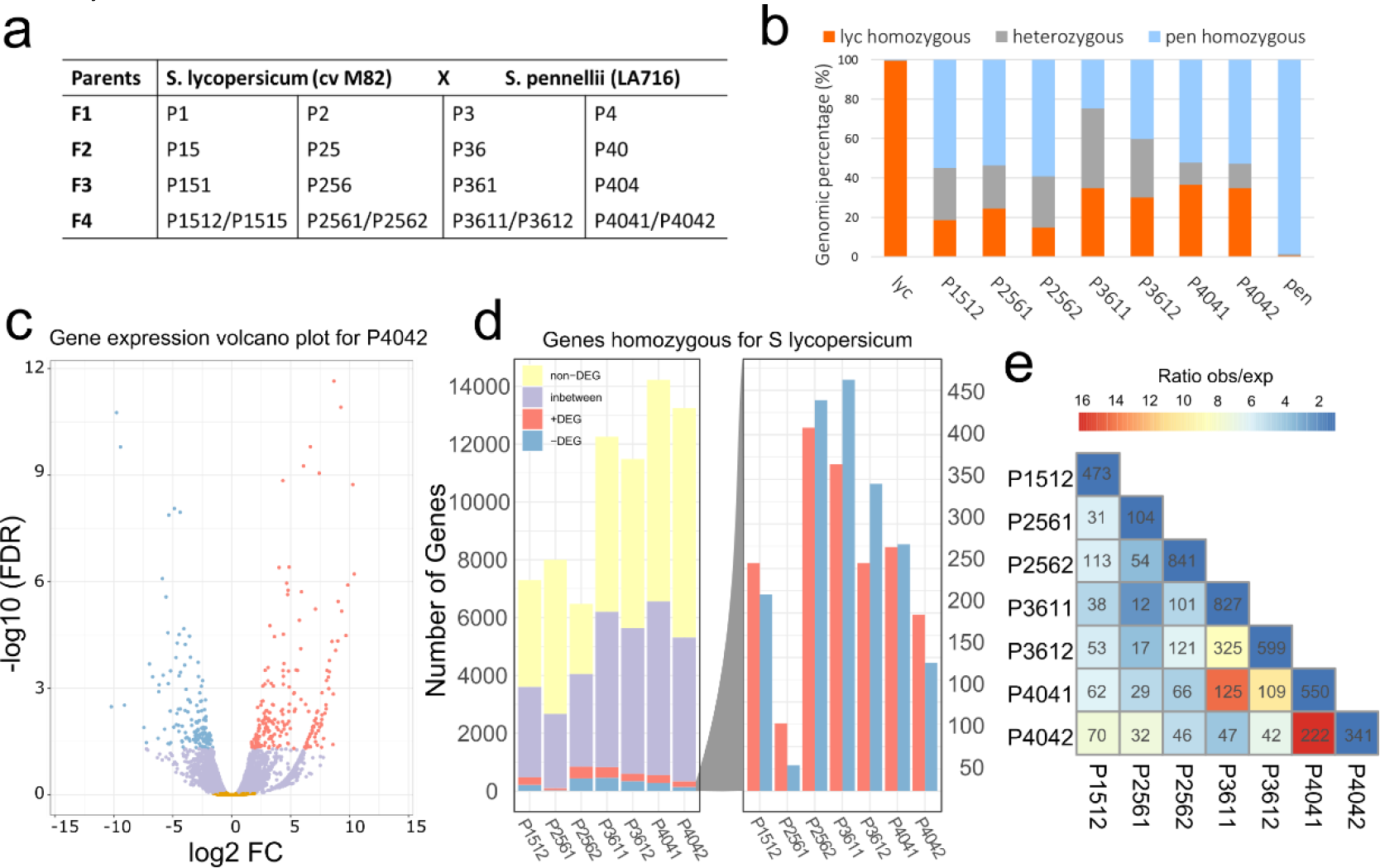
Gene expression analysis of *lyc* homozygous regions in hybrids. a. Parental lines *(S. lycopersicum* and *S. pennellii*) and offspring (from F1 to F4), each column being a separate F1 lineage. For F2-F4, each plant keeps the number of the parental plant adding one digit. The number of digits indicates the generation (i.e. P4042 is F4 generation, progeny of plant P404). b. Percentage of homozygous regions for *S. lycopersicum* (orange*), S. pennellii* (blue) or heterozygous (grey) for each of the F4 plants and parental lines. c. Volcano plot showing the distribution of genes upregulated (+DEG, red), downregulated (-DEG, blue), “in between” (lilac) and not differentially expressed (non-DEG, yellow) in *lyc* homozygous regions for P4042. Thresholds: DEG, FDR <0.05; “in between” genes, 0.05 <FDR < 0.9; non-DEG, FDR > 0.9. d. Distribution of number of *lyc* genes in each group for the F4s, being each bar an individual plant. The right-hand panel shows an expanded view of the +DEG (red) and - DEG (blue). e. Correlation table showing the number of *lyc* DEG genes shared among the F4 hybrids. Color scale shows the ratio of observed over expected genes shared in a pairwise combination. Ratio above 1 means the outcome is higher than expected.

To simplify bioinformatic analysis and avoid mapping biases we focused on expressed genes and sRNA loci (SL) in homozygous regions. (Fig.1b, ExtDataFig.2a). The general trend in seven F4 plants from four different lineages was that homozygosity in the F4s is greater for *pen* (average 48%) than for *lyc* (average 28%) and that heterozygosity (average 24%) is greater than anticipated (12.5%) (Fig.1b).

## Patterns of gene expression in hybrids deviated from those in the parents

From seven F4 RNAseq datasets we identified up- or down-regulated genes (+DEG and -DEG respectively (FDR<0.05)) (Fig.1c,d, ExtDataFig.3a) relative to the parents. There were also genes that were not differentially expressed (non-DEG (FDR>0.9)) and an ‘in between’ class that we could not assign to any of the other three classes with high confidence.

There were 6000 to 14000 *lyc* homozygous genes in each of the F4 datasets of which 1.2 to 12.6% were differentially expressed and equally divided between those that were up- or down-regulated (Fig.1d). There were similar patterns and numbers of differentially expressed genes from the *pen* homozygous regions (ExtDataFig.3a). The sets of differentially expressed *lyc* genes were most similar between sibling F4 plants, but there was also more overlap in differential expression than expected by chance in the four independent lineages (Fig.1e). From this observation we conclude that hybridisation induced mechanisms affecting gene expression are non-random: a differentially expressed gene in one plant is also likely to be similarly affected in independent lineages.

## sRNA loci from endogenous viruses were affected in hybrids

McClintock proposed that transposons would influence genomic modification due to the ‘shock’ of hybridisation promoting movement of transposable elements leading to structural rearrangement of the genome^1^. It is also possible that transposon-derived sRNAs could mediate altered patterns of gene expression in hybrids^4,5^. To test this latter possibility, we characterised sRNA profiles in the parental and hybrid lines.

The sRNA loci (SL) were distributed widely and, as with the gene expression profiles, they were divided in four classes based on whether they were up- or down-regulated (+DESL and -DESL), not differentially expressed (non DESL) or ‘in between’ (Fig.2a, ExtDataFig.3b). The differentially expressed SL accounted for 1.5-2.5% of the total (FDR<0.05) corresponding to thousands of loci. More of these loci were up-rather than down-regulated, consistent with hybridization triggering sRNA silencing that persists into the F4. As with the differentially expressed genes, the hybridisation-induced changes to SLs were most similar between sibling F4 plants but there were also more coincident patterns of differential expression than would be expected by chance in the independent lineages (Fig.2b): a differentially expressed SL in one plant was also likely to be similarly affected in independent lineages.

**Figure 2.**
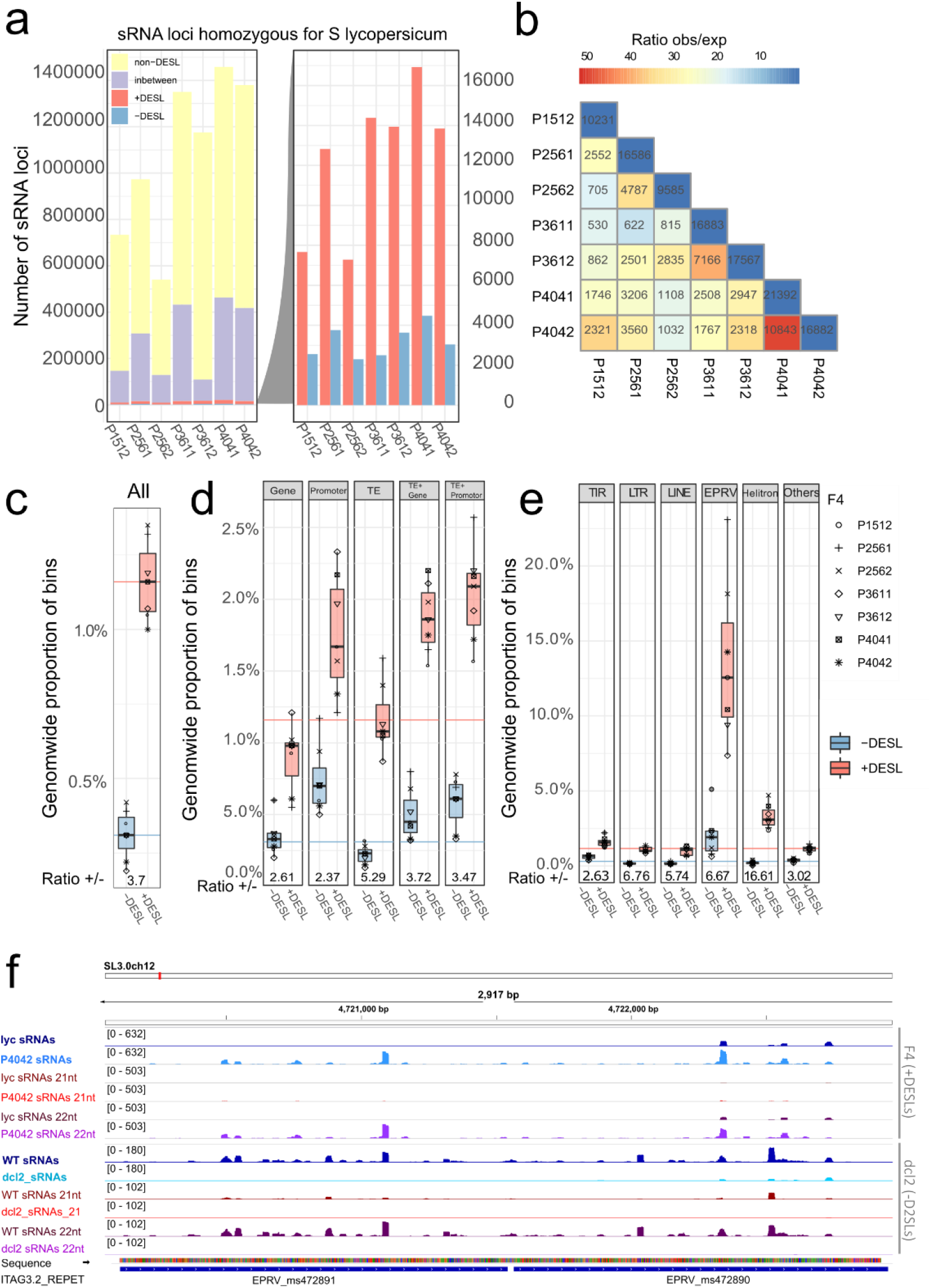
sRNA analysis of *lyc* homozygous regions in hybrids: a. *S. lycopersicum* genome is divided in 200nt bins (sRNA loci). Distribution of *lyc* sRNA loci divided in upregulated (+DESL, red), downregulated (-DESL, blue), “in between” (lilac) and non-differentially expressed (non-DESL, yellow). The right-hand panel shows an expanded view of the +DESL (red) and -DESL (blue). Thresholds: DESL, FDR <0.05; “in between”, 0.05 <FDR < 0.9; non-DESL, FDR > 0.9. b. Correlation table showing the number of *lyc* DESL shared among the F4 hybrids. Color scale shows the ratio of observed vs expected sRNA loci shared in a pairwise combination. Ratio above 1 means the outcome is higher than expected. c-e. Box plot showing the percentage of *lyc* sRNA loci that are +DESL or -DESL mapping (c) genome wide, (d) to each genomic feature, (e) to TE order. Ratio of +DESL/-DESL is shown at the bottom. Each plant is represented by different dot shapes. Box plots elements: box limits, upper and lower quartiles; center line, median; whiskers, from each quartile to the minimum or maximum. For comparison, colored lines are the percentage values for +DESL (red) and -DESL (blue) mapping the whole genome (from c). f. Genome browser snapshot of two EPRV elements (bottom) showing sRNA mapping (blue) and filtered for 21 nt (red) and 22nt (purple) sRNAs, for *lyc* and P4042 (top tracks); and WT and *dcl2* (bottom tracks). Numbers in brackets show the scale (normalized number of reads) for each track.

High level genome features (genes, promoters, transposable elements) overlapped similarly with differentially expressed *lyc* SLs (*lyc* DESLs, Fig.2c,d) and, for each of them, there were more up-rather than down-regulated loci, following the genome-wide pattern (Fig2.c). There is a similar profile of differentially expressed SLs in the *pen* genome although the bias to up-regulation was less pronounced than in *lyc* (ExtDataFig.3b-d).

However, analysis of transposable element types revealed that endogenous pararetroviruses (EPRV) and helitron SLs deviated from the genome-wide pattern. An exceptionally high proportion of SLs overlapping EPRVs and helitrons are differentially expressed (about 13% and 3%, respectively) (Fig.2e,f). For EPRV elements there was overrepresentation of both up- and down-regulated SLs whereas, for helitrons, the deviation from the genome-wide pattern was specifically with up-regulated loci. In the *pen* genome there was a similar over-representation of differentially expressed SLs amongst the EPRVs (ExtDataFig.3e). Genomic alignment of the sRNA sequence data implicates subsets of the EPRV elements in this differential expression, but they do not belong to a specific clade of the phylogeny tree (ExtDataFig.4).

## Differential expression of DCL2-dependent sRNA loci in hybrids

Small RNA size classes (21, 22 or 24nt) are associated with distinct RNA silencing pathways^6^. Most of the non-differentially expressed sRNAs were from the 24nt class (non-DESL, Fig.3a, ExtDataFig.5a) whereas the differentially expressed SLs especially those in the in the up-regulated class (+DESL, Fig.3a, ExtDataFig.5a) included 21nt and 22nt species. The 22nt sRNAs were predominant in differentially expressed SL mapping to TIR and LTR transposons and, strikingly, to EPRVs (Fig.2f, Fig.3a, ExtDataFig.5b).

**Figure 3.**
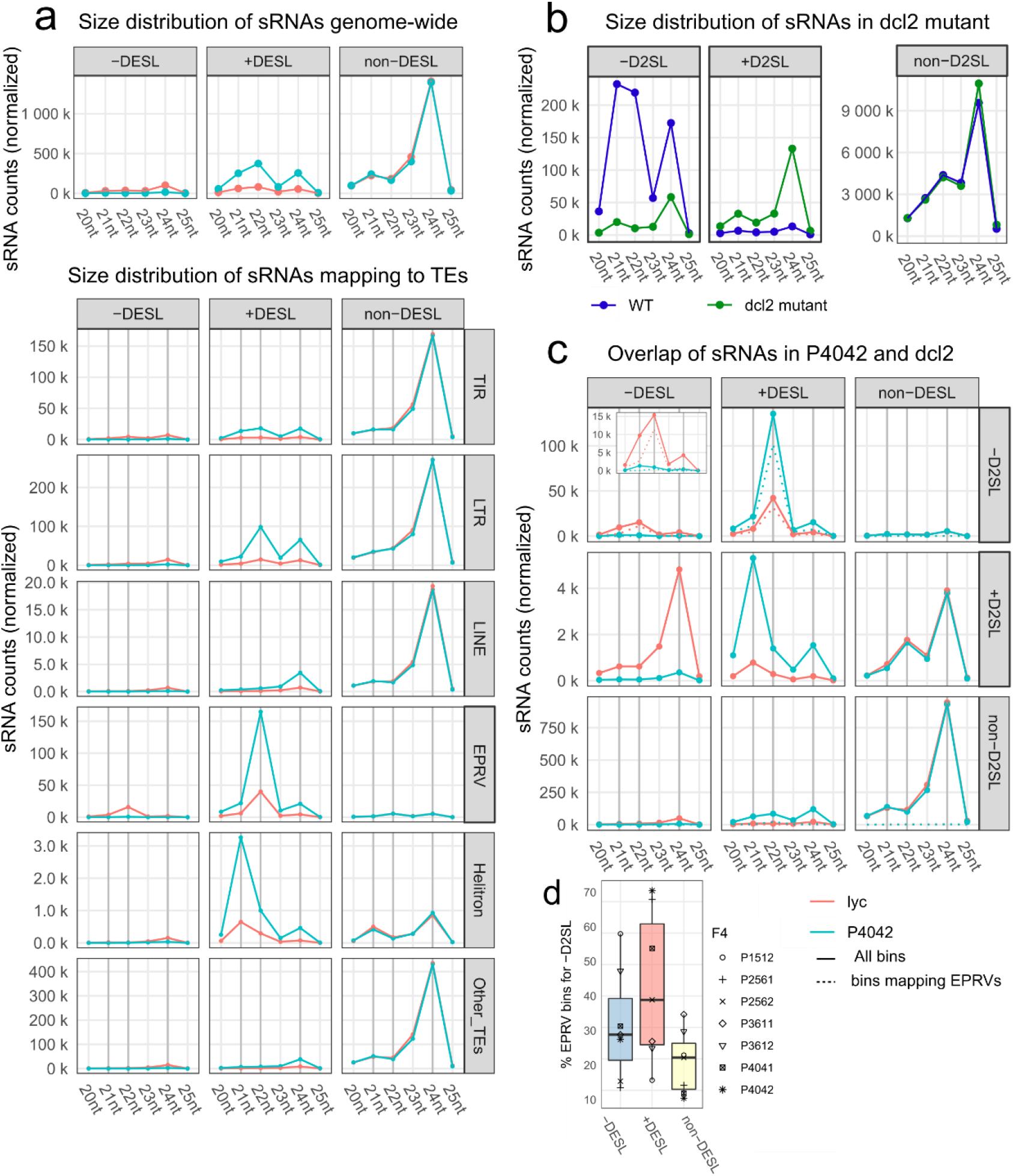
sRNA size distribution and DCL2 dependency. a. Normalized sRNA counts for P4042 (blue line) compared to lyc (red line) divided by size for -DESL, +DESL and non-DESL (top panel) and mapping to each TE order (bottom panels). b Normalised sRNA counts divided by size of sRNA loci in *dcl2* mutant (green) compared to its WT (purple), for -D2SL (downregulated sRNA loci), +D2SL (upregulated sRNA loci) and non-D2SL (non-differentially expressed sRNA loci). c. Normalised sRNA counts divided by size of sRNA loci in P4042 (-DESL, +DESL and non-DESL) overlapping with sRNA loci in *dcl2* (-D2SL, +D2SL and non-D2SL). Red line- sRNA counts for *lyc*, blue line -sRNA counts for P4042. Solid line- counts for all sRNA loci in each group category for that plant. Dotted line – counts for sRNAs in each category that map to EPRVs. Inset with expanded y axes for -DESL/-D2SL. d. EPRV contribution to each group on Fig3.c top panel: -D2SL-DESL (sRNA loci downregulated in F4s and *dcl2*), - D2SL+DESL (sRNA loci downregulated *dcl2*, upregulated in F4) and -D2SL, non-DESL (sRNA loci downregulated in *dcl2* with no change in F4, for comparison). Each plant is represented by different dot shapes. Box plots elements: box limits, upper and lower quartiles; center line, median; whiskers, from each quartile to the minimum or maximum.

The predominant mechanism for 22nt sRNA production involves DCL2^7,8^ and, correspondingly, the sRNA loci in WT tomato that decrease in *dcl2* (-D2SL) have abundant 21nt and 22nt species whereas those that increase (+D2SL) or are not affected (non-D2SL) are predominantly 24nt (Fig.3b). DCL2 clearly plays a major role in the F4 plants because sRNAs mapping to the loci that increase or decrease (+ or -DESL respectively) also map predominantly to the loci that are dependent on DCL2 (-D2SL) with the 22nt size class being prominent in all lines (Fig.3c, ExtDataFig.5c). Of these DCL2-dependent sRNAs from the SL that are up- or down-regulated in the F4 plants, a high proportion map to EPRVs (Fig.2f, Fig.3d, dotted line in + and -DESL/-D2SL panels of Fig.3c and ExtDataFig.5c).

This pattern of DCL2 dependency (-D2SL), 22nt abundance and EPRV representation was specific to the sRNAs that increased or decreased in the F4 plants (from + or – DESL) (Fig.3c). The sRNAs that were not affected in the hybrids (non-DESL) were also not dependent on DCL2 (non-D2SL) and were predominantly 24nt. Those that were changed in the hybrids (+ or -DESL) and increased rather than decreased in *dcl2* (+D2SL) were generally of lower abundance than the DCL2-dependent loci (-D2SL) and they were a mixture of 21 and 24nt species (Fig.3c, ExtDataFig.5c). These various patterns indicate, therefore, that the up- or down-regulated sRNAs in the F4 hybrids (+ and – DESL) are produced predominantly by DCL2 and they correspond to EPRV loci.

To find out whether the changes to sRNA expression occurred immediately after the *lyc* x *pen* hybridization or whether it was progressive over several generations, we analysed differentially expressed SL from the *lyc* parent and the F1-F4 generations (ExtDataFig.6). In each instance there was a shift to either up- or down-regulated SLs in the F1. For the up-regulated loci the change progressed into the F4 generation but for the down-regulated loci there was little further decrease in expression after the F1. It is likely therefore that the *lyc* x *pen* hybridization triggered changes to the SLs in the F1 and that there was a consequent change to the sRNA profiles that progressed further in the F2-F4 generations.

## DCL2 affects gene expression in *lyc* x *pen* hybrids

Associated with the sRNA changes in *dcl2* there were also up- and down-regulated genes (+D2G and - D2G respectively)^8^. If there is a causal link involving DCL2, 22nt sRNAs and changes in gene expression in the *lyc* x *pen* hybrids, we predicted an overlap of the genes that are differentially expressed in the F4s (DEG) and in *dcl2* (D2G) relative to the parental or wild type plants. To test this prediction we reanalysed the published *dcl2* and WT RNAseq datasets^8^ and compared the differentially expressed genes with the up- and down-regulated genes in our F4 hybrids.

This analysis (Fig.4a,b, ExtDataFig.7a) revealed an extraordinary coincidence in the patterns of gene expression affected by *dcl2* and the *lyc x pen* hybridisation both in terms of the gene identity and the quantitative effect on the expression level. The coincident identity was reflected in 23% of the genes upregulated in *dcl2* (+D2G) also increasing in the F4s (+DEG) and 27% of the downregulated genes in *dcl2* (-D2Gs) decreasing in the F4s (-DEGs). These values were highly non-random: only 3% of the genes upregulated in *dcl2* (+D2Gs), for example, overlapped with downregulated genes in the F4s (-DEGs) and 3% of the -D2Gs coincided with +DEGs. The correlated quantitative effects were reflected in the near-linear relationship in the level of up- or down-regulation in the F4s and *dcl2* (Fig.4b, ExtDataFig.7b). From these patterns we conclude that the hybridisation of *lyc* x *pen* triggered a change to the DCL2-mediated pathway of sRNA production that influenced the profile of gene expression in the F4 progeny.

**Figure 4.**
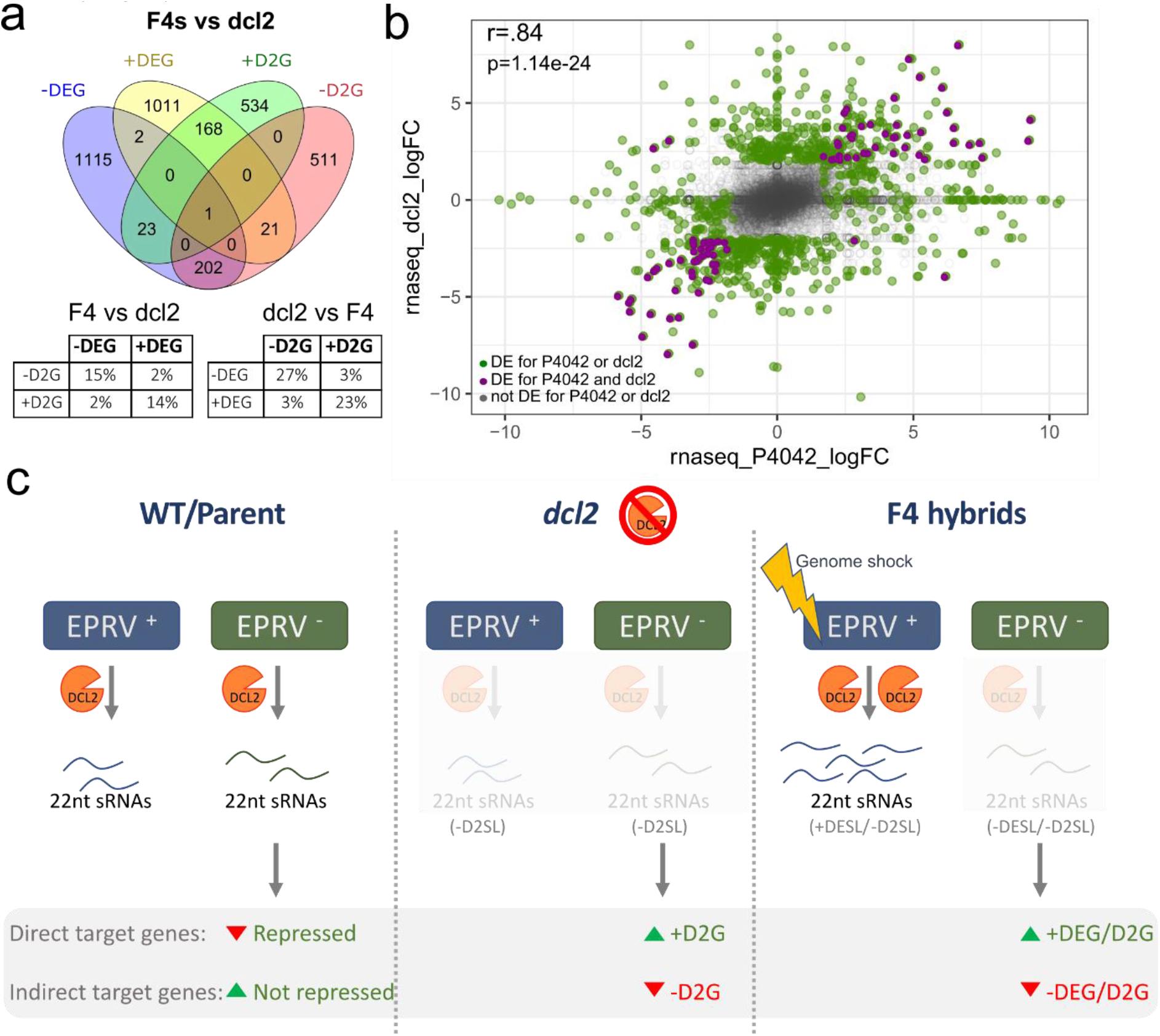
Common response in the change of gene expression in the F4 hybrids and *dcl2* mutant. a. Venn diagram of the consolidated DEG for the F4s (compilation of +/−DEGs in at least one F4 plant) compared to differentially expressed genes in *dcl2* (+/−D2G). Tables showing the percentage of DEGs in the F4 (being 100%) shared with *dcl2* (left), and for DE genes in *dcl2* (being 100%) shared with F4 (right). b. Scatter plot of the fold change (FC) in gene expression in P4042 vs *dcl2*. Grey: non-DE. Green: DE genes for P4042 or *dcl2*. Purple: DE genes for both P4042 and *dcl2*. Pearson correlation coefficient value (r) for DEG and D2G genes (purple genes), p=p-value. c. Model of the contribution of EPRVs, 22nts and DCL2 to hybrid genetic profile: in the *dcl2* mutant, as DCL2 is not present, transcripts from EPRV^+^ and EPRV^−^ loci are not processed into sRNAs (-D2SL), producing upregulation of genes (+D2G) that were repressed in WT. In F4s, the upregulation of EPRV+ leads to production of 22nt sRNAs. There are different possibilities for the competition between these pathways, one option is that in the F4s, although DCL2 is present at normal levels, it is required to process the upregulated EPRV+ transcripts (+DESL/−D2SL). Thus, DCL2 may not be available to process transcripts from EPRV^−^ locus (-DESL/-D2SL) leading to gene upregulation (+DEG/D2G). This leads to a change in the sRNA population, beyond those produced from EPRVs, with similarities between *dcl2* and the F4s (+DESL/+D2SL) that may be responsible for the change in gene expression of indirect targets (-DEG/-D2G).

## Discussion

Through these analyses of sRNAs we show that DCL2, 22nt sRNAs and EPRV elements are implicated in up- and down-regulation of gene expression following the hybridisation of *lyc* and *pen*. Perturbation of the sRNA profile initiated in the F1 plants following the hybridisation event (ExtDataFig.6) and it persisted in later generations until at least the F4 (Fig.2, ExtDataFig.6 and ExtDataFig.3b). EPRV-specific 22nt sRNAs were key components of the modified sRNA profile (Fig. 3, ExtDataFig.5b) and, as in a *dcl*2 mutant, there was a consequent effect on gene expression (Fig.1, Fig.4). Other sRNA changes involving helitrons and LTR elements also occurred, but they were less prominent than those involving EPRVs (Fig.2e).

The overlapping changes to gene expression in *lyc* x *pen* hybrids and the *dcl2* mutant were not due to suppression of DCL2 because the F4 hybrids had DCL2 levels similar to WT and they had abundant 22nt sRNAs. An alternative explanation implicates the DCL2-dependent sRNA that were up or down-regulated in the F4 progeny (-D2SL and + or – DESL in Fig. 3c, ExtDataFig.5c). Most of these sRNAs were from EPRV loci and, in our model (Fig. 4c), we refer to those down-regulated in the F4 as EPRV^−^ and up-regulated as EPRV^+^. We hypothesise that a hybridisation-induced genome shock led to the increased production of sRNAs from the EPRV^+^ loci.

In the wild type/parental plants the EPRV^−^ sRNAs could have down-regulated gene expression by directly targeting mRNAs and they could have up-regulated genes indirectly if the direct targets were suppressors of gene expression. In the *dcl2* mutant the EPRV^−^ sRNAs were reduced or absent (Fig. 3c, ExtDataFig.5c) so that the down-regulated genes in the WT would have been up-regulated (+D2G) and the up-regulated WT genes would have been down-regulated (-D2Gs) (Fig.4c).

In *lyc* x *pen* hybrids we envision that genome shock-induced activation of the EPRV^+^ loci suppressed the sRNA production from the EPRV^−^ loci due to competition for limiting amounts of DCL2 or if the EPRV^+^ sRNAs silenced the long RNA precursors of the EPRV^−^ sRNAs (Fig.4c). In either scenario the EPRV^−^ pathway would have been suppressed and, as observed, the effects on gene expression would have been the same as in the *dcl2* mutant (+DEG ≡ +D2G and -DEG ≡ -D2G, Fig.4a,b, ExtDataFig.7).

These mechanisms affecting gene expression in the *Solanum* hybrids differ from the genome shock in other organisms^1,3,9^ in that they involve endogenous viruses (EPRVs) rather than transposons. The EPRV inserts are likely fragments of viral genomes that infected ancestors of the modern *Solanum* species^10,11^ but, nevertheless, they have retained features that link them with the DCL2 antiviral RNA silencing pathway^7^.

This genome shock is also different from P element-mediated hybrid dysgenesis in *Drosophila*^2,3^, inter-specific rice hybrids^9^ and the various examples described by McClintock^1^ in which the consequence was transposon mobilization leading, in some instances, to chromosomal restructuring. In the *Solanum* example described here the perturbation of the DCL2-mediated RNA silencing is a less drastic effect. It leads to both up- and down-regulation of gene expression in the hybrid and there could be both positive and negative phenotypic effects, as observed (ExtDataFig.1a,b). This sRNA-based mechanism involving EPRVs in other species or transposons could modify gene expression in hybrid populations and increase the range of traits available for selection in natural evolution or breeding for agriculture.

With the involvement of 22nt sRNAs these RNA silencing effects could be more complex and far-reaching than with other size classes of sRNA. The 22nt size class is specifically associated with translational regulation^12^ and it also has the potential to trigger secondary sRNAs that target multiple mRNAs^13,14^. There is potential for similar outcomes in the many other species carrying EPRVs including bitter orange, rice, lucky bamboo, Dahlia, pineapple, grapes, poplar and fig^10^. In sugar beet^15^ and soybean^16^ the EPRV are associated with 22nt sRNAs as in *Solanum*. This sRNA-based mechanism involving EPRVs could modify gene expression in hybrid populations of many species and increase the range of traits available for selection in natural evolution or breeding for agriculture.

## METHODS

### Plant material

Parental lines (Tomato (*Solanum lycopersicum*) cv. M82 and *Solanum pennellii* LA716) and hybrid population derived from the crosses were grown from seeds in compost (Levington M3) and maintained in the Botanical Garden Greenhouses (24-18°C, 16 hr/8 hr day/night regime) propagated by cuttings so that hybrids and parents could be sampled simultaneously. A replica of the collection was maintained at NIAB (Cambridge, UK). Samples for library preparation were collected for parent and progeny the same day in a period of 2h. Each sample is a pool of 3-5 young leaves (1 to 3cm). Sample replicates for each plant were collected with at least one week of difference. To compare the plants along the generations for the phenotypic analysis, we grew 4 cuttings per plant for 8 weeks. Dry weight was calculated by weighting the areal part of each plant after being dried in an oven at 65C for 1 week.

### RNA extraction and library preparation (mRNAseq and sRNAseq)

RNAseq: 5 μg of total Trizol-purified RNA purified was treated with Turbo DNase free kit (Thermo Fisher), ribosomal depleted using Ribo-Zero Kit (Epicentre) and libraries prepared using ScriptSeq v2 RNA-Seq Library Preparation Kit (Epicentre), PCR amplified in 9 cycles. In total, 12 libraries were prepared, containing two biological replicates for each of the plant. Equimolecular pooled libraries were sequenced with the Illumina High Output Kit v2 150PE in a NextSeq 550 benchtop machine (Illumina) at the SLCU (UK).

sRNAseq: 1 μg of total Trizol-purified RNA purified was used for library preparation using the NEBNext Multiplex Small RNA Library Prep Set for Illumina (NEB) with libraries indexed in 12 cycles PCR. Size selection was performed using BluePippin 3% agarose cassettes (Sage Science). Equimolecular pooled libraries were sequenced with the Illumina High Output Kit v2 30 SE in a NextSeq 550 benchtop machine (Illumina) at the SLCU (UK).

### Reference genome and annotation

Reference genome consists of a merged genome where genome assemblies for both parental strains, *Solanum Pennellii*^17^ and *Solanum lycopersicum* cv. Heinz^18^ assembly version SL3.0 were combined into one reference, also including mitochondrial and chloroplast genomes. The merged genome was sliced into 200bp adjacent non-overlapping bins. Bins were used as intervals as a basis to count various features which either overlap (such as gene or TE annotation) or map into them (e.g. sRNAs/RNA-seq reads). Gene annotation and transposable elements are based ITAG3.2 (“gene_models” and REPET_repeats_aggressive”) (https://solgenomics.net). MicroRNA annotation was obtained from miRBase release 22.1^19^. Since precursor coordinates were only available for Heinz assembly version 2.50, we performed a lift over to Heinz assembly 3.0 using Liftoff version 1.6.1^20^.

### Genotyping using RNA-Seq reads

To genotype each F4, we used the SNP information in the RNAseq reads for actively transcribed regions. To determine SNPs that differentiate between the 2 parents, we performed a whole genome alignment using Mummer v3.23^21^. As first step the subprogram “nucmer” with default settings was used to perform a genome wide alignment between both assemblies. The delta results were then filtered with “delta-filter -r -q” to exclude chance and repeat induced alignments, leaving only the “best” alignments between both assemblies (only SNPs on the *S. lycopersicum* genome that exhibit homologous alignments from *S. pennellii* and vice versa), resulting in 180868 alignment blocks. Those were used as basis to extract SNPs positions using “show-snps –CTr” resulting in 23,528,724 SNPs which positions were available for both assemblies.

Mapped RNA-Seq reads (mapping was performed only on one assembly at a time to ensure that homologous genes map to the same coordinates to allow genotyping) were processed using “bcftools mpileup -d10000 -Ov -R $snps -o ${bam}.R.vcf -f $ref ${bam}.bam” for all sequencing libraries to generate RNA-Seq genotype information. The SNP positions (focusing on the *S. lycopersicum* genome) were then used to determine the genotype for each of SNP by comparing the RNA-Seq derived SNPs with the expected SNP derived from the mummer analysis above using the R package “vcfR” and custom scripts. For each mummer SNP, if RNA-Seq information supported only the expected *S. lycopersicum / S. pennellii* genotype this SNP would be annotated as *S. lycopersicum / S. pennellii* homozygous respectively. If both variants occurring, it was annotated as heterozygous. Since this process can be error-prone and we expected the same genotype over larger regions we constructed a hidden Markov model (HMM) using the R package “HMM” to construct a genome wide genotype map. The assumption was that genotype patterns are relatively large (typically mega bases) containing multiple regions with transcriptional activity, i.e. the genotype of an interval between to transcriptional location of the same genotype is likely to be of the same genotype. Since this process is subject to noise and a genotype sections is expected to be large, the HMM Viterbi algorithm was parameterized to account for this using 3 states (homozygous *S. lycopersicum*, homozygous *S. pennellii* and heterozygous) with respective probabilities (0.45, 0.45, 0.1). Transition probabilities between states were chosen to be very small (1e-9). This map also allows to infer the genotype of non RNA-Seq covered sections.

### Small RNA sequencing data processing and analysis

Small RNA libraries were computationally processed using the pipeline available at https://github.com/seb-mueller/snakemake_sRNAseq. Briefly, small RNAs reads (fastq format) were subjected to 3’ adaptor removal (trimming) using cutadapt v3.1 removing Illumina universal adapters. Sequences with <15nt and >40nt in length were discarded, and the remaining sequences were mapped to the reference genome. Specifically, mapping was performed using Bowtie version 1.2 with 2 different approaches. Firstly, keeping multi mapping reads with parameters “bowtie --wrapper basic-0 -v 0 -k 1 -m 50 --best -q” which randomly assigns reads to a mapping location among the multiple locations and secondly only keeping uniquely mapping reads with “bowtie --wrapper basic-0 -v 0 -k 1 -m 1 --best –q”. Both were performed requiring 0 mismatches (-v 0). For both approaches, mapped sRNAs reads were counted separately for each library based on overlapping bins as defined above. An additional read counting was also performed for each small RNA size class (20-25nt length) for each bin for a more detailed analysis.

Differential expression (DE) analysis was performed using the R/bioconductor package edgeR with first conduction ‘glmFit’ to fit genewise negative binomial followed by ‘glmLRT’ which conducts likelihood ratio tests. Test intervals were bins as and TMM normalization method was used for ` calcNormFactors` library normalisation. Comparisons were performed between parents and F4 libraries separately. Multiple test correction was carried calculating the false discovery rate (FDR) from p-values using the Benjamini & Hochberg method as implemented in the R function “p.adjust”.

Bins were grouped based on FDR threshold into non-DESL (FDR > 0.9), (0.05 < FDR < 0.9) and DESL (differentially expressed small RNA locus; FDR < 0.05). DESL were further classed into +DESL (F4 > WT) and –DESL (F4 < WT). Bins with insufficient coverage (edgeR default) were excluded from statistical analysis and assigned “none” class.

### RNA sequencing processing and analysis

RNA-Seq libraries were trimmed using cutadapt v3.1 and mapped using STAR version 2.7.5c using “-- outFilterMultimapNmax 20 --alignSJoverhangMin 8 --alignIntronMin 20 --alignIntronMax 10000 -- bamRemoveDuplicatesType UniqueIdentical --outFilterMismatchNmax 20” as parameters against the merged reference genome described above. Quantification of gene expression quantification was performed using the R/bioconductor “Rsubread” package. Differential expression (DE) analysis was performed using the R/bioconductor package edgeR using genes as testing intervals and TMM normalization method. Comparisons were performed between parents and F4 libraries separately. Multiple test correction was carried calculating the false discovery rate (FDR) from p-values using the Benjamini & Hochberg method as implemented in the R function “p.adjust”. Genes were grouped based on FDR threshold into non-DEG (FDR > 0.9), (0.05 < FDR < 0.9) and DEG (differentially expressed smallRNA locus; FDR < 0.05). DESL were further classed into +DESL (F4 > WT) and –DESL (F4 < WT). Genes with insufficient coverage (edgeR default) were excluded from statistical analysis and assigned “none” class.

### External *dcl2* library re-analysis

sRNA-Seq: Raw sRNAs libraries^7^ were downloaded from SRA WT=SRR6436051, SRR6436050 and *dcl2*_mutant=SRR6436048, SRR6436055) and pre-processed the same way as described for the our sRNA-Seq libraries. The adapter sequence used for trimming was found to be “AGATCGGAAGAGCAC” (Illumina adapter). WT libraries were compared against *dcl2* mutant libraries using the same threshold as our sRNA analysis. Similarly, bins were classified into -D2SL for *dcl2* < WT and +D2SL for *dcl2* > WT. RNA-Seq: Raw sRNAs libraries^8^ were downloaded from SRA WT=SRR6866906, SRR6866908, SRR6866909 and *dcl2*_mutant=SRR6868335, SRR6868336, SRR6868333 and preprocessed as described previously. WT libraries were compared against *dcl2* mutant libraries using the same threshold as our RNAseq analysis. Similarly, genes were classified into -D2G for *dcl2* < WT and +D2G for *dcl2* > WT.

### Phylogenetic analysis

In order to perform an unbiased phylogenetic analysis on EPRV domains for both *S. lycopersicum* and *S. pennellii*, we retrieved consensus nucleic acid sequences encoding for env, inclusion body, movement protein, and polymerase from GIRI repbase (LycEPRV_I) and aligned them to both genome assemblies using BLAT version 36 using with following parameters “-minScore=25-minIdentity=80 - noHead-maxIntron=1000”. For each domain BLAT alignments were imported into R using “hiReadsProcessor” library and mapping location were overlapped (based on genomic location) with available annotations as well as bins which come with their respective annotations (again for both genomes). The underlying genomic sequences were written into fasta files for each location encoding annotation in the naming header. All sequences found for a specific domain (env, inclusion body, movement protein, and polymerase) were multiple aligned with “t_coffee” version 13.41.0.28bdc39 using default parameters.

Resulting multiple alignments were subjected to trimAl version 1.4 rev15 to remove gaps (usging “-gappyout“), spurious sequences or poorly aligned regions. Cleaned up alignments were used to build a phylogentic tree using RAxML-NG version 1.0.1, a phylogenetic tree inference tool which uses maximum-likelihood (ML) optimality criterion (“raxml-ng --msa $aln --model GTR+G“).”). Phylogenetic tree visualization of resulting trees was performed using FigTree version 1.4.4.

### Data visualisation and statistical analysis

Plots were carried out using ggplot2^22^. Statistical tests were carried out in R version 4.0. Pearson correlation was calculated using “cor.test” R function with default parameters (two sided).

### Data and material availability

Biological material is available by request to the corresponding author. Raw data is available in ArrayExpress repository. smallRNAseq (sRNAseq data from the F4s and parental lines): E-MTAB-10613; RNAseq (RNAseq data from the F4s and parental lines): E-MTAB-10660.

### Code availability

Computationally reproducible scripts are available at https://github.com/seb-mueller/scripts_tribe

## Abbreviations

SL: sRNA loci
EPRV: endogenous pararetroviruses
DEG: differentially expressed genes in F4s compared to parental lines
D2G: differentially expressed genes in *dcl2* compared to WT
DESL: Differentially expressed sRNA loci in F4s compared to parental lines
D2SL: Differentially expressed sRNA loci in *dcl2* compared to WT

## Acknowledgements

The authors thank Melanie Steer, Antonia Yarur, CUBG and NIAB staff for horticultural and technical assistance; Dr Baster for assistance with library sequencing; Prof Jiggins and Ian Warren for providing access to Blue Pippin; SLCU for providing access to NextSeq550; and Prof Henderson, Dr Harris, Dr Martinho, Dr Singh, Dr Deng and Dr Putra for helpful comments on the manuscript. This work was supported by European Research Council Advanced Investigator grant ERC-2013-AdG 340642 (Transgressive Inheritance in plant Breeding and Evolution [TRIBE]), the Royal Society (RP170001), the Balzan Foundation and the Broodbank Fund. SLG is a Senior Broodbank Research Fellow. D.C.B. is the Royal Society Edward Penley Abraham Research Professor.

## Author contributions

D.C.B., S.L.G. and S.Y.M designed the study; S.L.G. performed the biological experiments; S.Y.M. and S.L.G. performed the data analysis; D.C.B., S.L.G. and S.Y.M. wrote the manuscript.

## Competing interests

The authors declare no competing interests.

## EXTENDED DATA LEGENDS

**Extended data Figure 1. Phenotyping variability of the hybrid population along the generations.** For comparison, 4 cuttings per plant were grown for 8 weeks. **a.** The 5th leaf from the bottom was used for the comparative phenotypic analysis. All pictures at the same scale being scale bar 10cm. **b.** Quantitative analysis of phenotypical traits (dry weight, height, and number of leaves). Parental lines in red, F1-F3 plants in light blue and F4 plants in dark blue. Dotted lines show the values for the parental lines for comparison to identify transgenic phenotypes. Box plots elements: box limits, upper and lower quartiles; center line, median; whiskers, 1.5x interquartile range; points, outliers.

**Extended data Figure 2. F4 genomes are chimeras of *S. lycopersicum* and *S. pennellii*.** RNAseq-based SNP genotyping for each of the F4 plants (P1511, P2561, P2562, P3611, P3612, P4041, P4042). Top track (within each chromosome panel): For each tomato nuclear chromosome is it shown the genotypic results (red: region homozygous for S lycopersicum, green: heterozygous, light blue: homozygous for S. pennellii, pink: could not be determined). Bottom track: chromatin status divided into euchromatin (light grey) or heterochromatin (dark grey) as described in Wang, Z. and Baulcombe, D.C. Nat Comm (2020) 11:1221. As expected, there were very few meiotic crossovers in the pericentromeric region. Preferential inheritance of some regions of the genomes including those that were homozygous *lyc* in all F4 plants (regions in chromosomes 2,8,10 and 11) may due to selection against genes or combinations of genes that caused failure to produce progeny in some F2 and F3 progeny.

**Extended data Figure 3. Gene expression and sRNA analysis for the S pennellii homozygous regions for each F4. a.** Distribution of number of genes in each group for the F4s, being each bar an individual plant. The right-hand panel shows an expanded view of the +DEG (red) and -DEG (blue). **b.** Distribution of sRNA loci divided in upregulated (+DESL, red), downregulated (-DESL, blue), “in between” (lilac) and non-differentially expressed (non-DESL, yellow). The right-hand panel shows an expanded view of the +DESL (red) and -DESL (blue). Thresholds: DESL, FDR <0.05; “in between” DESL, 0.05 <FDR < 0.9; non-DESL, FDR > 0.9. **c-e.** Box plot showing the percentage of *pen* sRNA loci that are +DESL or -DESL (**c**) genome wide, (**d**) to each genomic feature, (**e**) to TE order. Ratio of +DESL/-DESL is shown at the bottom. Each plant is represented by different dot shapes. Box plots elements: box limits, upper and lower quartiles; center line, median; whiskers, from each quartile to the minimum or maximum. For comparison, colored lines are the percentage values for +DESL (red) and -DESL(blue) mapping the whole genome (obtained from **c**).

**Extended data Figure 4. Phylogenetic relationship of EPRV envelope sequence.** Polar Tree layout of the phylogenetic tree build based on envelope sequences found on *S. lycopersicum* by manual BLAT search using as original the envelope sequence from EPRV on GIRI, followed by RaxML tree construction. Node names consist of domain name (envelope), followed by coordinates on genome (chromosome name and start coordinate), overlapping transposon family according to REPET annotation (NA if no overlap), and overlapping type of sRNA loci highlighting +DESL in red and -DESL in blue.

**Extended data Figure 5. Size distribution of sRNA loci in the F4s.** Name of plant shown at the top of each panel. **a.** Normalised sRNA counts divided by size for -DESL, +DESL and non-DESL and **b** mapping to each TE order. **c.** Normalised sRNA counts divided by size of sRNA loci in each F4 (-DESL, +DESL and non-DESL) overlapping with sRNA loci in *dcl2* (-D2SL, +D2SL and non-D2SL). Red line-sRNA counts for *lyc*, blue line -sRNA counts for each F4. Solid line- counts for all sRNA loci in each group category for that plant. Dotted line – counts for sRNA in each category that map to EPRVs.

**Extended data Figure 6. Heatmap of the inheritance of DESLs in each family.** Normalised reads for each generation for the sRNA loci identified as lyc DESL in one of the F4s per family (in bold letter on top), filtered for |FC| >9 and FDR < 1e-10. The strict threshold was used to reduce the size of the plot by focusing on the most significant changes. Normalization was performed for each bin by centering and scaling (z-score) across the lineages (rows). The names of bins (rows) correspond to the genomic coordinates of their first base. Annotation tracks were derived by overlapping the respective bins with genomic annotation features.

**Extended data Figure 7. Differentially expressed genes are shared between the F4s and *dcl2* mutant. a.** Venn diagram of genes upregulated (+DEG) and downregulated (-DEG) for an F4 plant compared to the genes upregulated (+D2G) or downregulated (-D2G) in the *dcl2* mutant. **b.** Scatter plot of the gene expression log fold change (FC) for each F4 compared to the *dcl2* mutant. Grey: non-DEG or in between for any of the two plants; green: DEG for the F4 or *dcl2*; purple: DEG for F4 and *dcl2*. Pearson Correlation coefficient values (r) for the DEG and D2G genes (purple genes) for each scatter plot, p = p value.

## Notes

### Competing Interest Statement

The authors have declared no competing interest.

