## Extended_data for "Endogenous virus sRNA regulates gene expression following genome shock in tomato hybrids"

**Supplementary Figures**

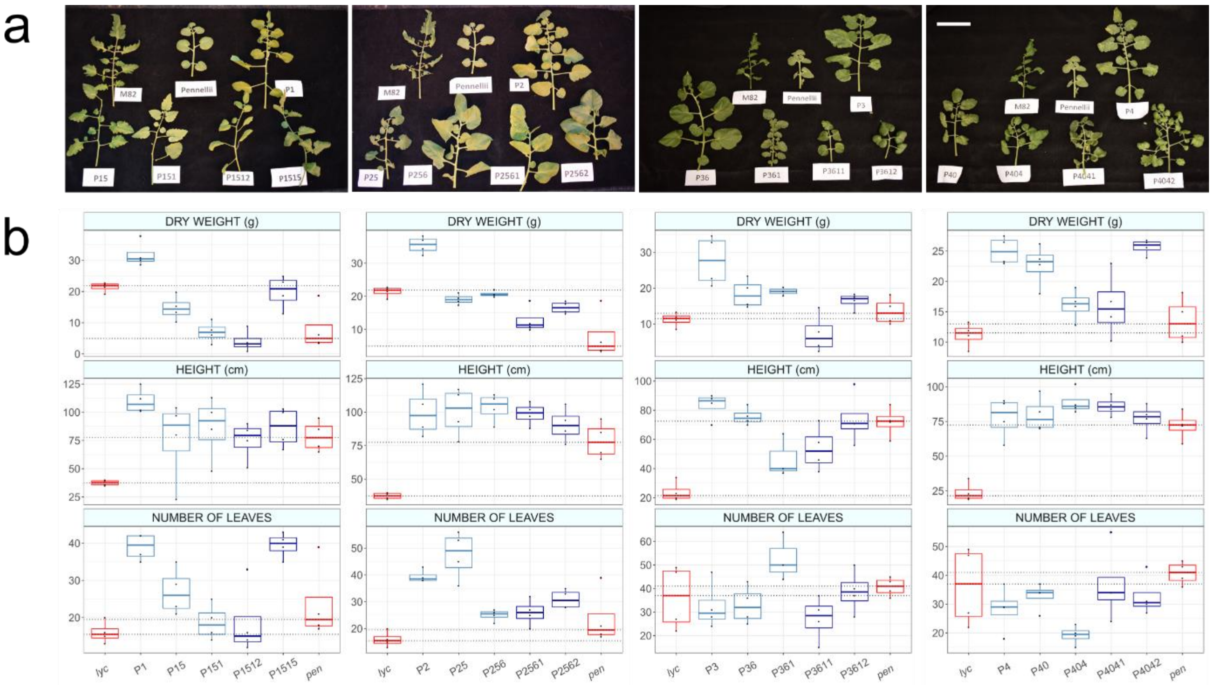


**Extended data Figure 1. Phenotyping variability of the hybrid population along the generations.** For comparison, 4 cuttings per plant were grown for 8 weeks**. a.** The 5th leaf from the bottom was used for the comparative phenotypic analysis. All pictures at the same scale being scale bar 10cm. **b.** Quantitative analysis of phenotypical traits (dry weight, height, and number of leaves). Parental lines in red, F1-F3 plants in light blue and F4 plants in dark blue. Dotted lines show the values for the parental lines for comparison to identify transgenic phenotypes. Box plots elements: box limits, upper and lower quartiles; center line, median; whiskers, 1.5x interquartile range; points, outliers.

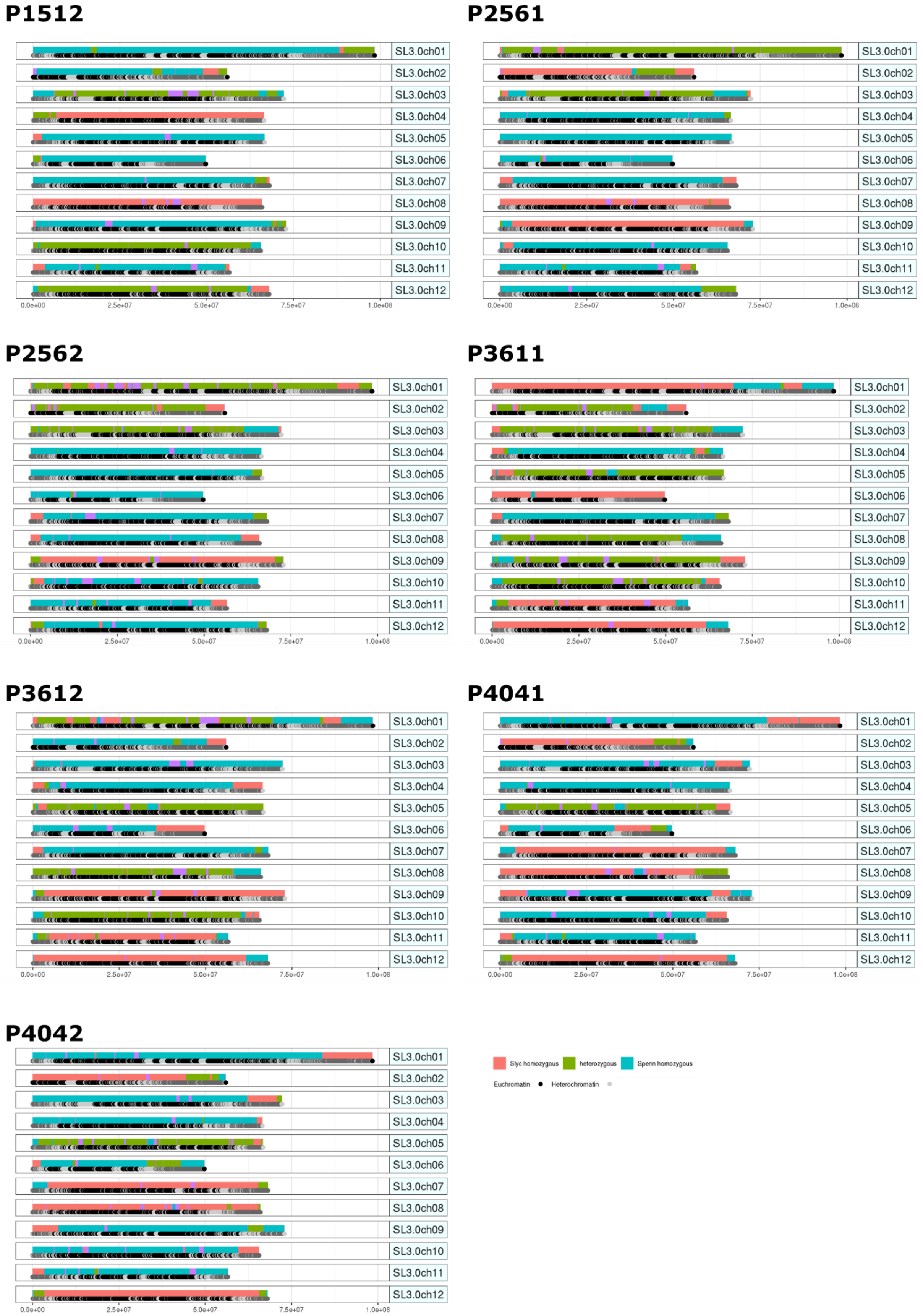


**Extended data Figure 2. F4 genomes are chimeras of *S. lycopersicum* and *S. pennellii*.** RNAseq-based SNP genotyping for each of the F4 plants (P1511, P2561, P2562, P3611, P3612, P4041, P4042). Top track (within each chromosome panel): For each tomato nuclear chromosome is it shown the genotypic results (red: region homozygous for S lycopersicum, green: heterozygous, light blue: homozygous for S. pennellii, pink: could not be determined). Bottom track: chromatin status divided into euchromatin (light grey) or heterochromatin (dark grey) as described in Wang, Z. and Baulcombe, D.C. Nat Comm (2020) 11:1221. As expected, there were very few meiotic crossovers in the pericentromeric region. Preferential inheritance of some regions of the genomes including those that were homozygous *lyc* in all F4 plants (regions in chromosomes 2,8,10 and 11) may due to selection against genes or combinations of genes that caused failure to produce progeny in some F2 and F3 progeny.

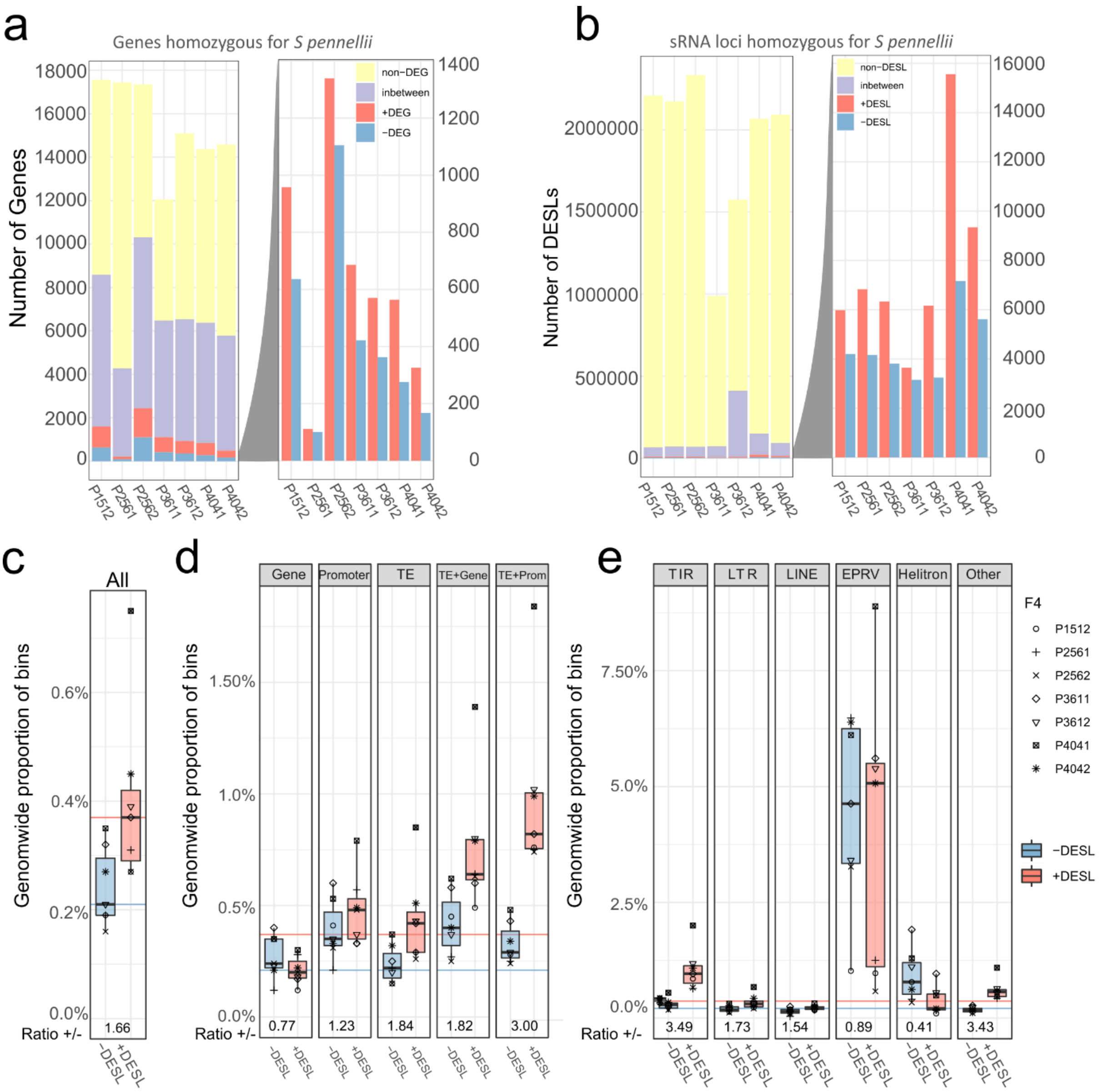


**Extended data Figure 3. Gene expression and sRNA analysis for the S pennellii homozygous regions for each F4. a.** Distribution of number of genes in each group for the F4s, being each bar an individual plant. The right-hand panel shows an expanded view of the +DEG (red) and -DEG (blue). **b.** Distribution of sRNA loci divided in upregulated (+DESL, red), downregulated (-DESL, blue), "in between" (lilac) and non-differentially expressed (non-DESL, yellow). The right-hand panel shows an expanded view of the +DESL (red) and -DESL (blue). Thresholds: DESL, FDR <0.05; "in between" DESL, 0.05 <FDR < 0.9; non-DESL, FDR > 0.9. **c-e.** Box plot showing the percentage of *pen* sRNA loci that are +DESL or -DESL (**c**) genome wide, (**d**) to each genomic feature, (**e**) to TE order. Ratio of +DESL/-DESL is shown at the bottom. Each plant is represented by different dot shapes. Box plots elements: box limits, upper and lower quartiles; center line, median; whiskers, from each quartile to the minimum or maximum. For comparison, colored lines are the percentage values for +DESL (red) and -DESL(blue) mapping the whole genome (obtained from **c**).

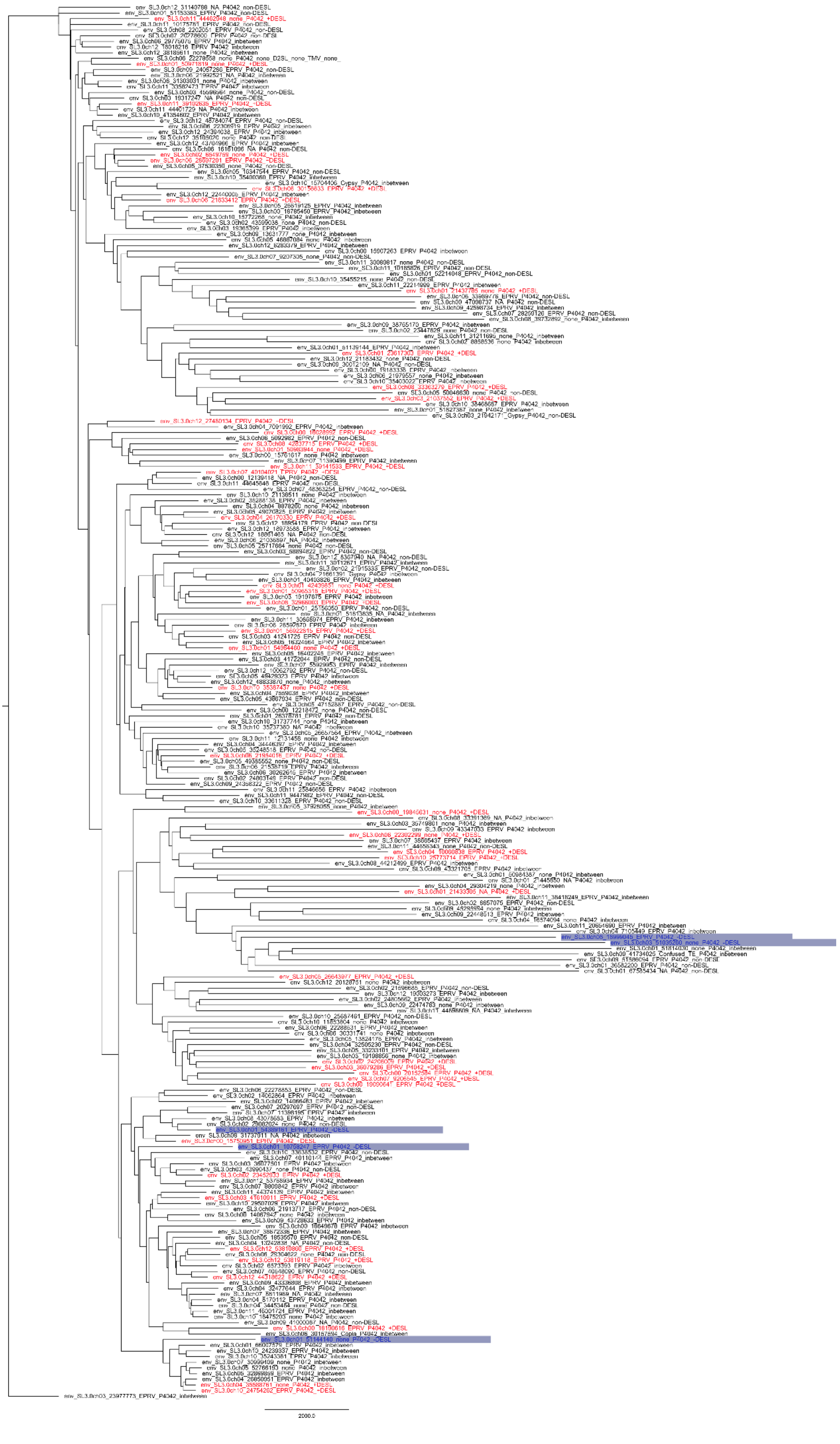
2000.0

**Extended data Figure 4. Phylogenetic relationship of EPRV envelope sequence.** Polar Tree layout of the phylogenetic tree build based on envelope sequences found on *S. lycopersicum* by manual BLAT search using as original the envelope sequence from EPRV on GIRI, followed by RaxML tree construction. Node names consist of domain name (envelope), followed by coordinates on genome (chromosome name and start coordinate), overlapping transposon family according to REPET annotation (NA if no overlap), and overlapping type of sRNA loci highlighting +DESL in red and -DESL in blue.

P1512

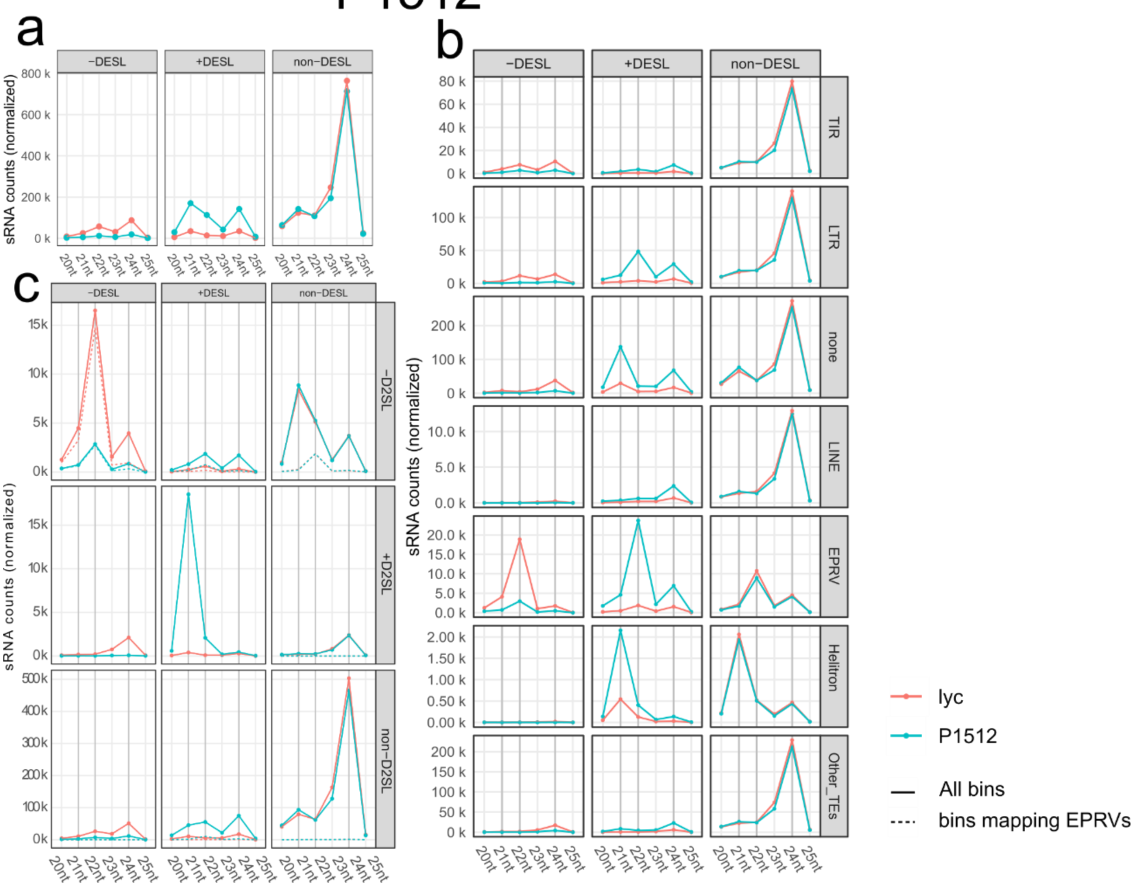
a
b
c
−DESL
+DESL
non−DESL
sRNA counts (normalized)
TIR
LTR
none
LINE
EPRV
Helitron
Other_TEs
−D2SL
+D2SL
non−D2SL
20nt
21nt
22nt
23nt
24nt
25nt
lyc
P1512
All bins
bins mapping EPRVs

P2561

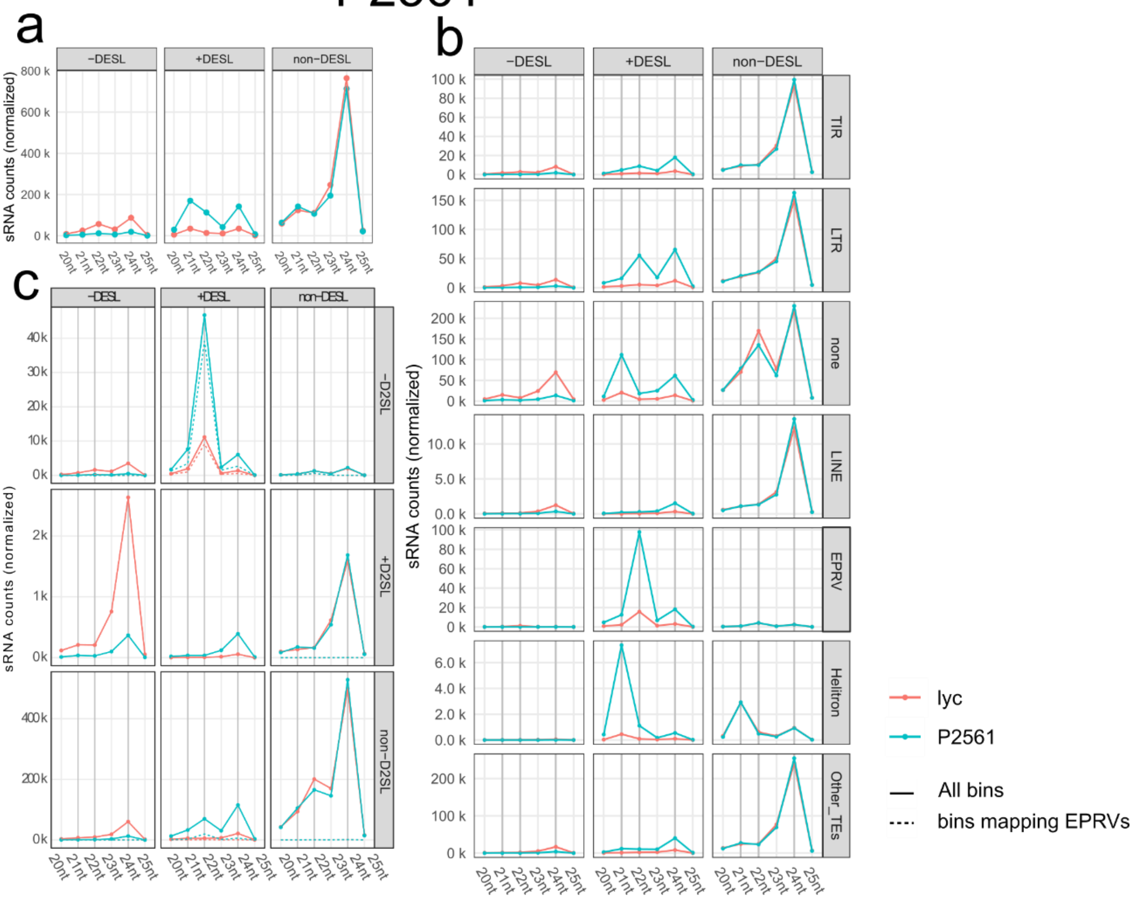
a
b
c
−DESL
+DESL
non−DESL
sRNA counts (normalized)
TIR
LTR
none
LINE
EPRV
Helitron
Other_TEs
−D2SL
+D2SL
non−D2SL
20nt
21nt
22nt
23nt
24nt
25nt
lyc
P2561
All bins
bins mapping EPRVs

# P2562

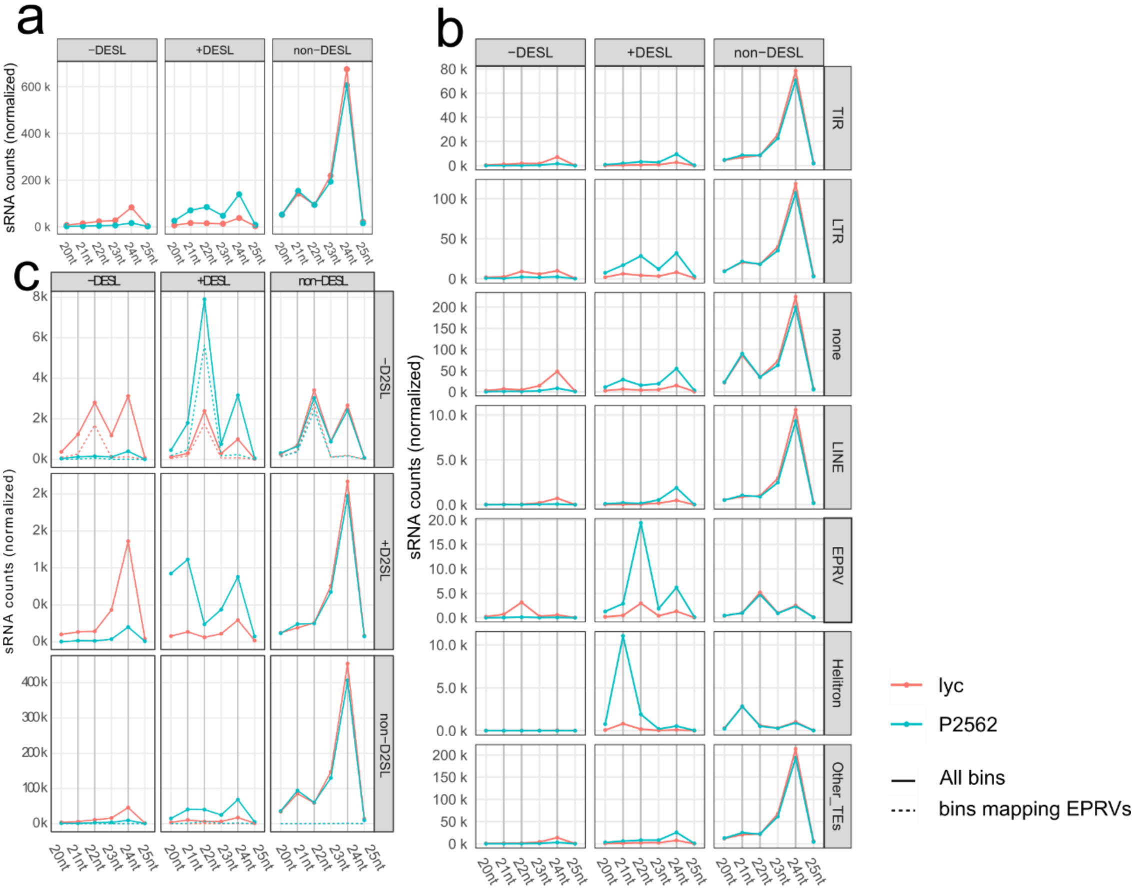
a
b
c
-DESL
+DESL
non-DESL
TIR
LTR
none
LINE
EPRV
Helitron
Other_TEs
-D2SL
+D2SL
non-D2SL
sRNA counts (normalized)
20nt
21nt
22nt
23nt
24nt
25nt
lyc
P2562
All bins
bins mapping EPRVs

# P3611

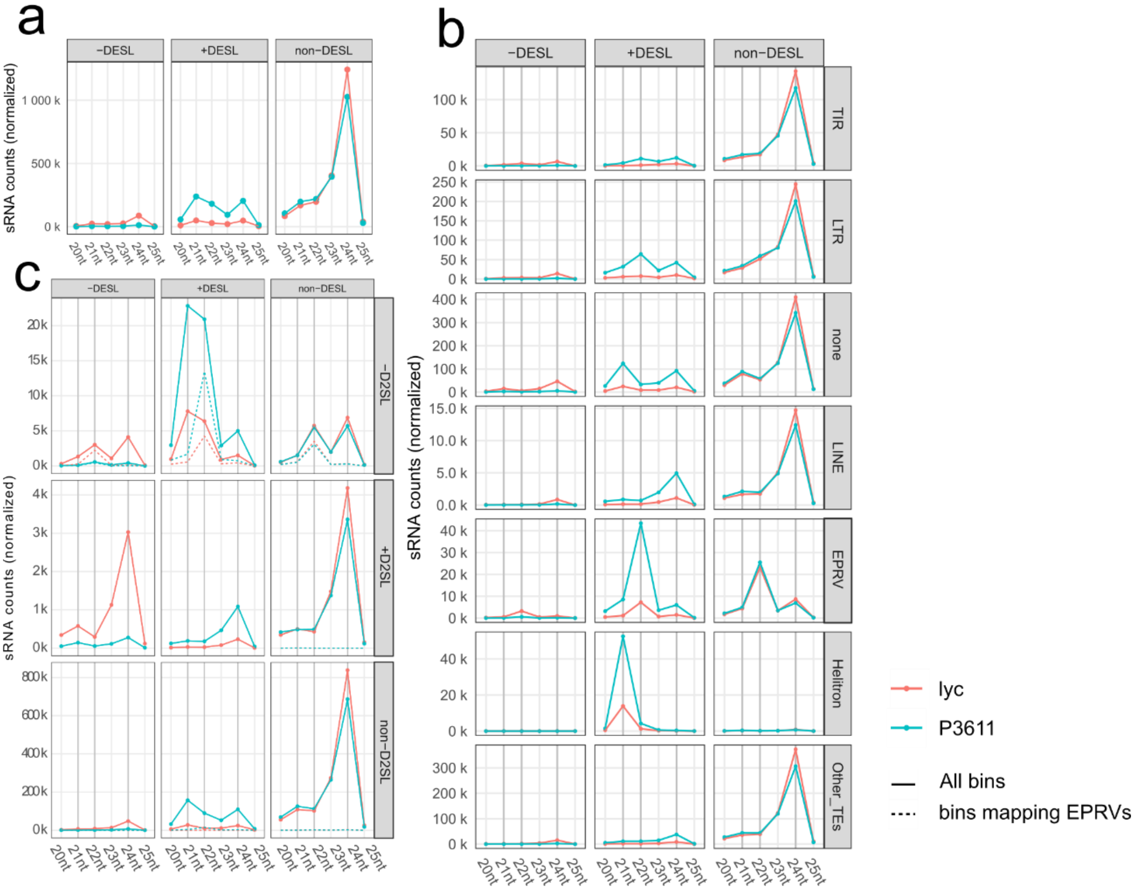
a
b
c
-DESL
+DESL
non-DESL
TIR
LTR
none
LINE
EPRV
Helitron
Other_TEs
-D2SL
+D2SL
non-D2SL
sRNA counts (normalized)
20nt
21nt
22nt
23nt
24nt
25nt
lyc
P3611
All bins
bins mapping EPRVs

# P3612

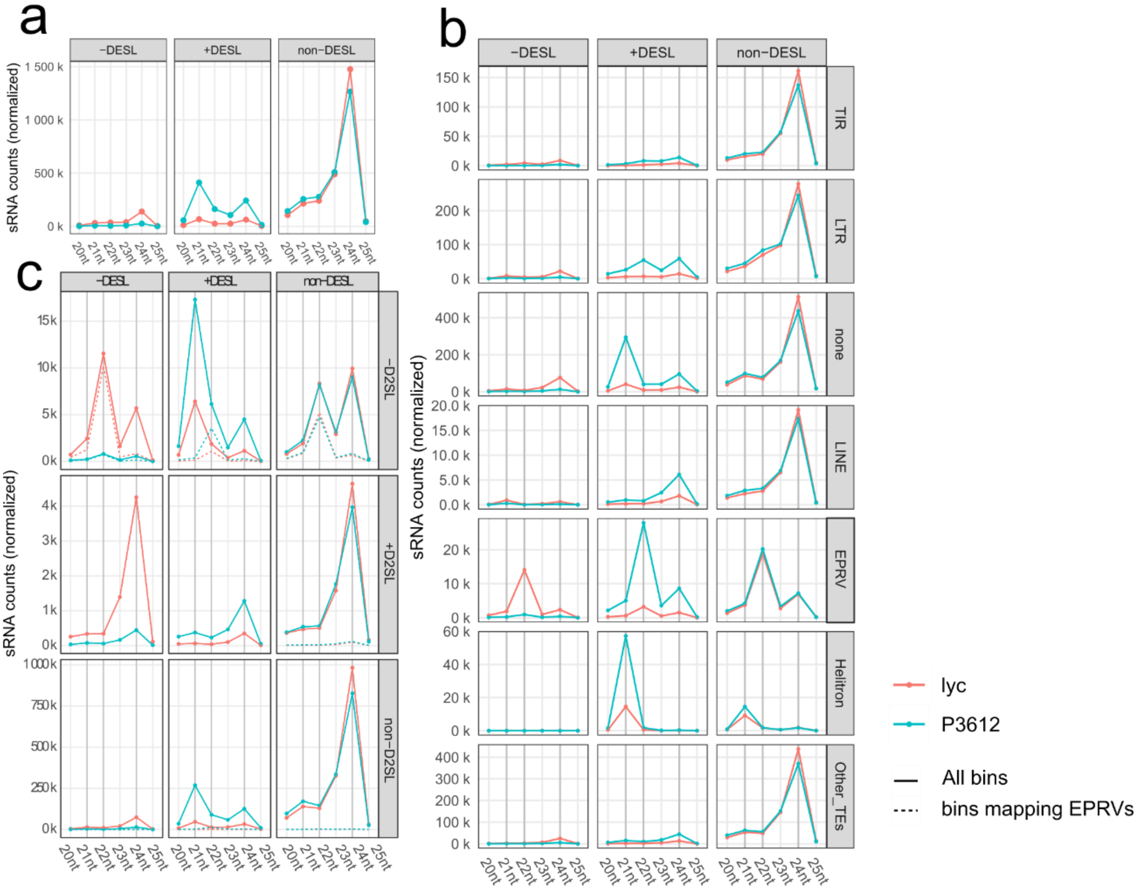

a
b
c
-DESL
+DESL
non-DESL
-D2SL
+D2SL
non-D2SL
TIR
LTR
none
LINE
EPRV
Helitron
Other_TEs
sRNA counts (normalized)
20nt
21nt
22nt
23nt
24nt
25nt
lyc
P3612
All bins
bins mapping EPRVs


# P4041

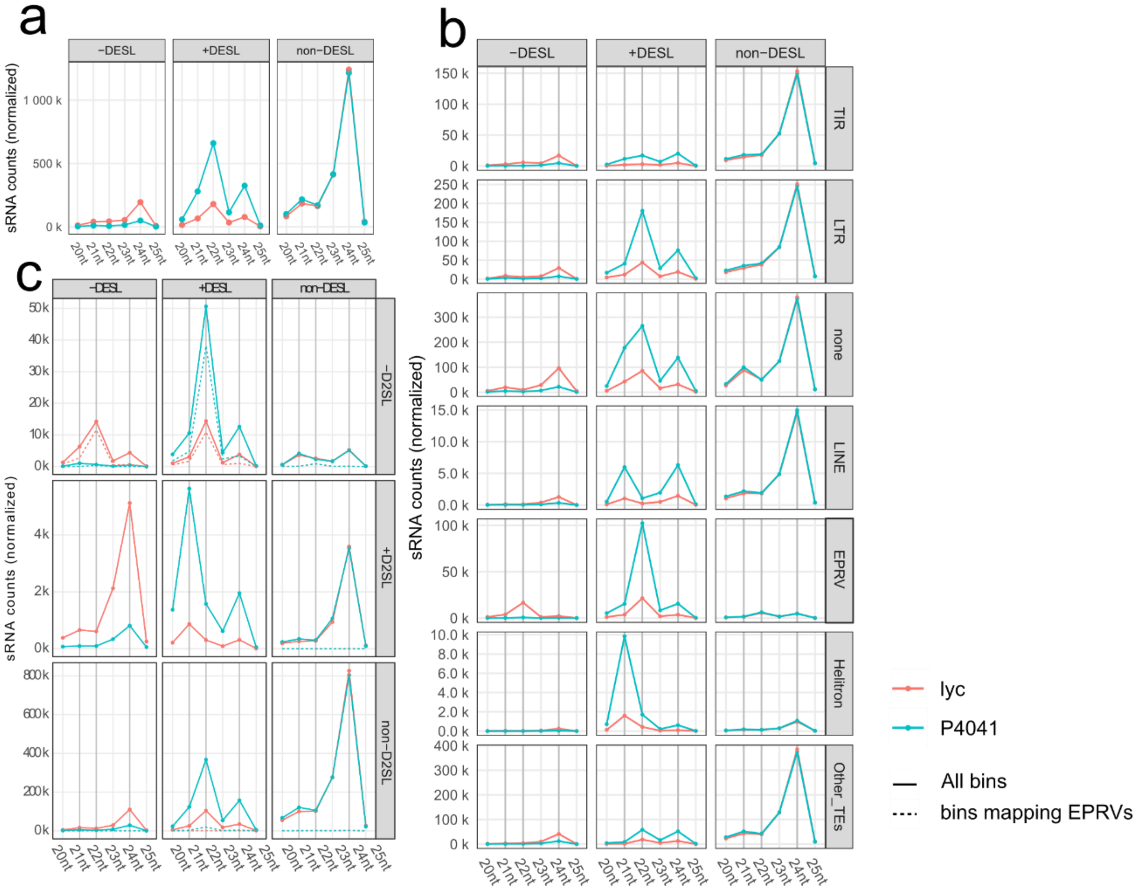

a
b
c
-DESL
+DESL
non-DESL
-D2SL
+D2SL
non-D2SL
TIR
LTR
none
LINE
EPRV
Helitron
Other_TEs
sRNA counts (normalized)
20nt
21nt
22nt
23nt
24nt
25nt
lyc
P4041
All bins
bins mapping EPRVs

**Extended data Figure 5. Size distribution of sRNA loci in the F4s.** Name of plant shown at the top of each panel. **a.** Normalised sRNA counts divided by size for -DESL, +DESL and non-DESL and **b** mapping to each TE order. **c.** Normalised sRNA counts divided by size of sRNA loci in each F4 (-DESL, +DESL and non-DESL) overlapping with sRNA loci in *dcl2* (-D2SL, +D2SL and non-D2SL). Red line- sRNA counts for *lyc*, blue line -sRNA counts for each F4. Solid line- counts for all sRNA loci in each group category for that plant. Dotted line – counts for sRNA in each category that map to EPRVs.

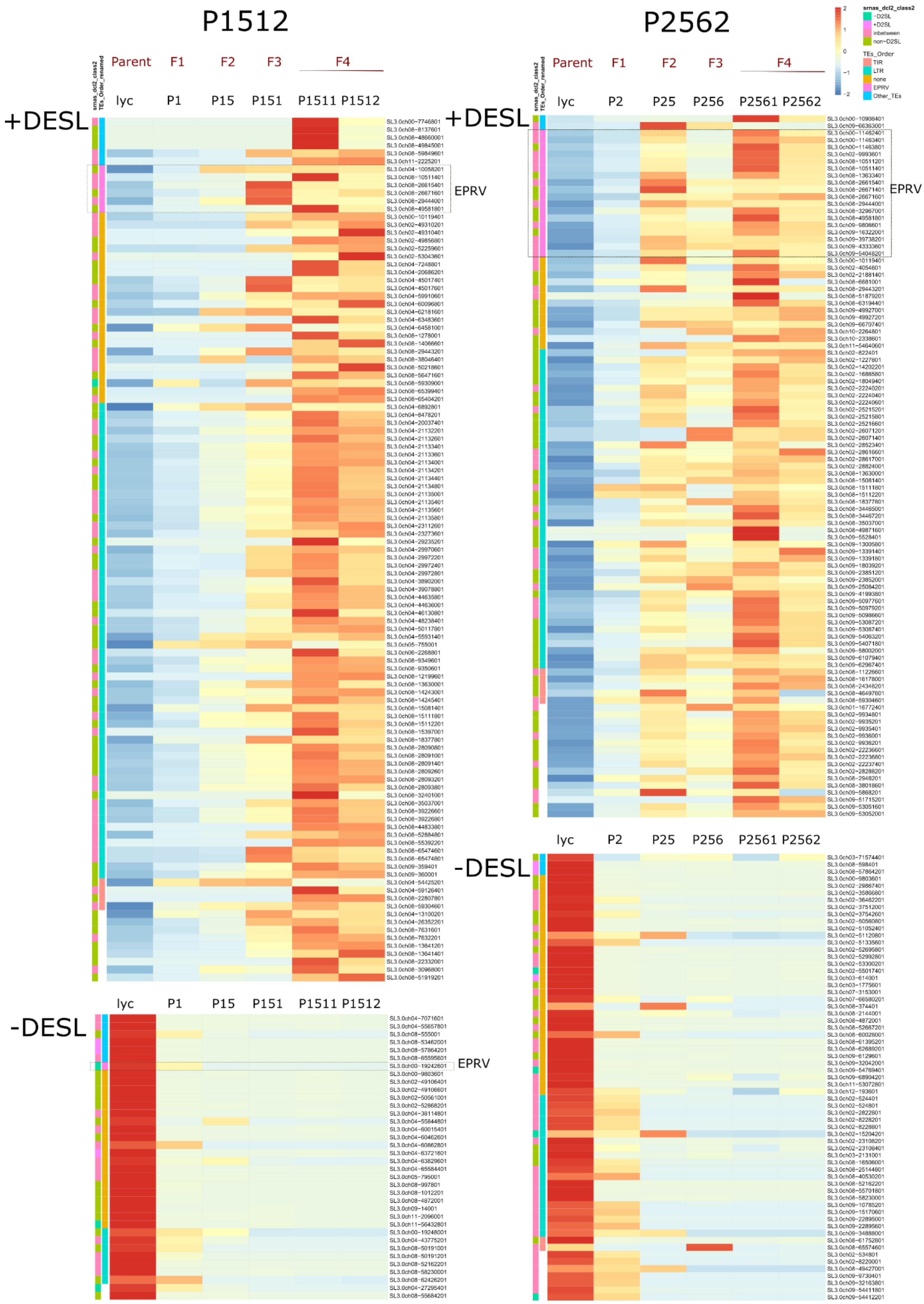
P1512
P2562
Parent
F1
F2
F3
F4
lyc
P1
P15
P151
P1511 P1512
P2
P25
P256
P2561 P2562
+DESL
-DESL
EPRV
srnas_dcl2_class2
TEs_Order_renamed
2
1
0
−1
−2
srnas_dcl2_class2
−D2SL
+D2SL
inbetween
non−D2SL
TEs_Order
TIR
LTR
none
EPRV
Other_TEs

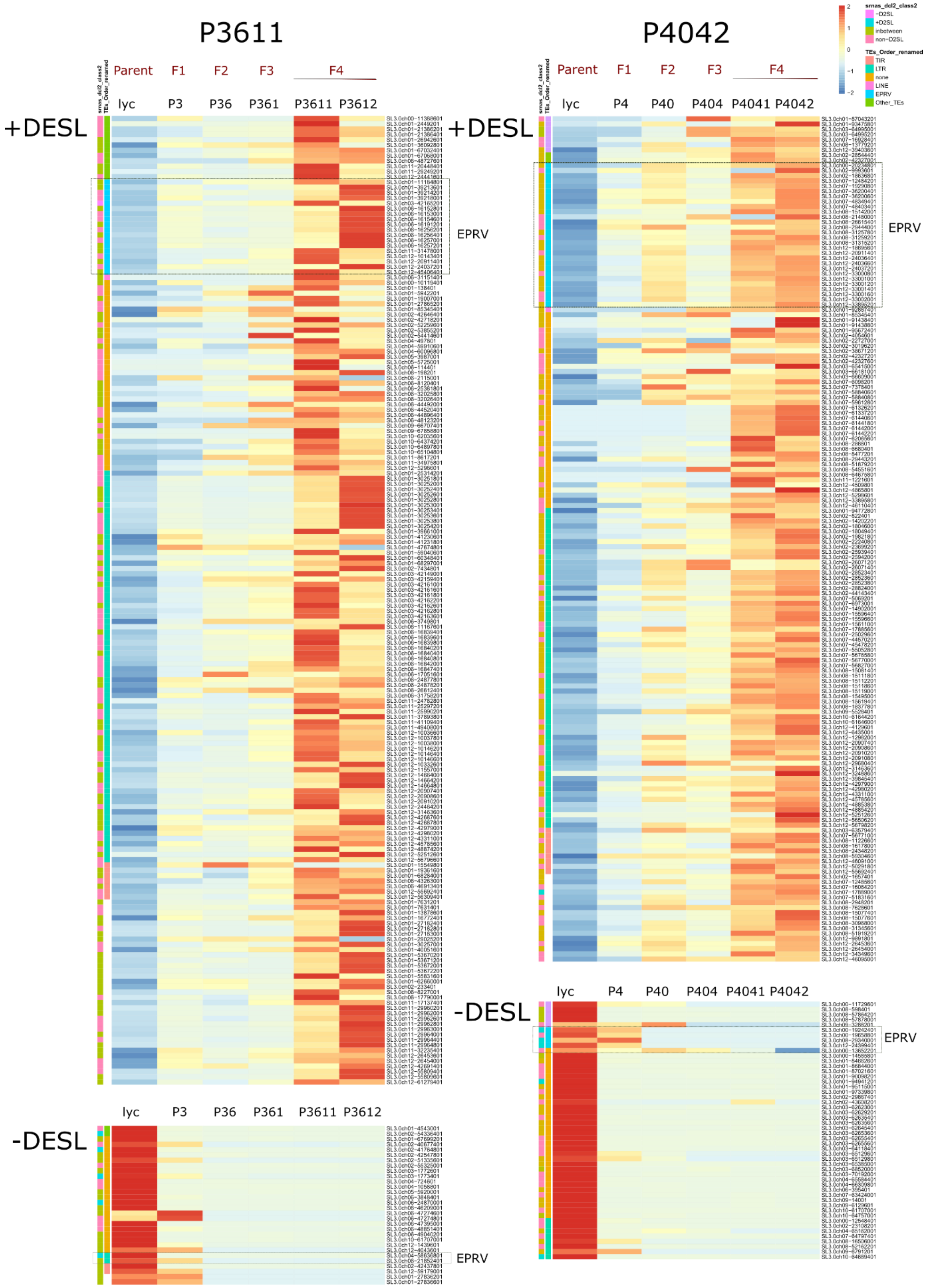
P3611
P4042
Parent
F1
F2
F3
F4
lyc
P3
P36
P361
P3611
P3612
P4
P40
P404
P4041
P4042
+DESL
-DESL
EPRV
srnas_dcl2_class2
-D2SL
+D2SL
inbetween
non-D2SL
TEs_Order_renamed
TIR
LTR
none
LINE
EPRV
Other_TEs

**Extended data Figure 6. Heatmap of the inheritance of DESLs in each family.** Normalised reads for each generation for the sRNA loci identified as lyc DESL in one of the F4s per family (in bold letter on top), filtered for |FC| >9 and FDR < 1e-10. The strict threshold was used to reduce the size of the plot by focusing on the most significant changes. Normalization was performed for each bin by centering and scaling (z-score) across the lineages (rows). The names of bins (rows) correspond to the genomic coordinates of their first base. Annotation tracks were derived by overlapping the respective bins with genomic annotation features.

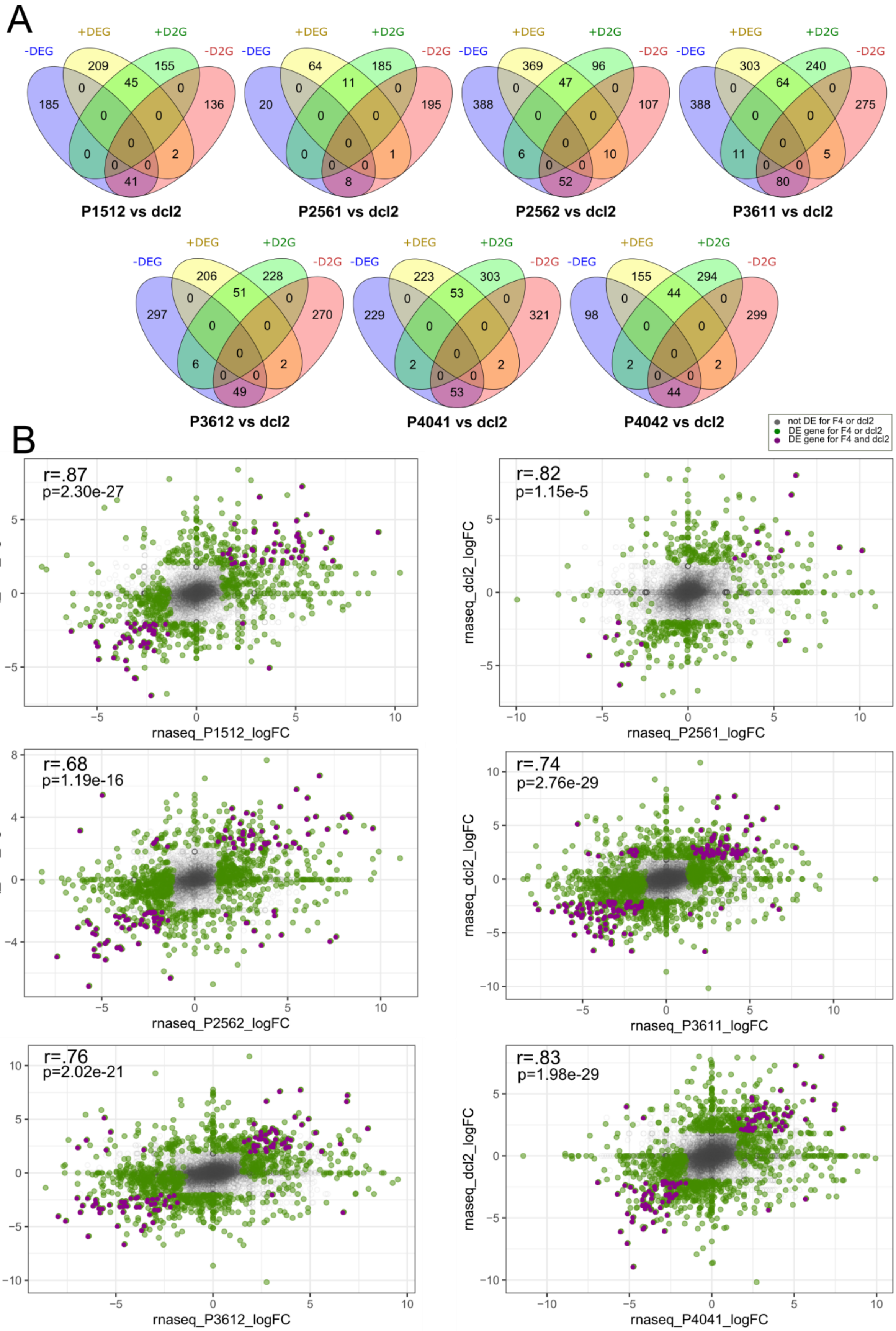

A
-DEG
+DEG
+D2G
-D2G
185 209 45 155 136 0 0 0 0 0 0 2 0 0 41
P1512 vs dcl2
20 64 11 185 195 0 0 0 0 0 0 1 0 0 8
P2561 vs dcl2
388 369 47 96 107 0 0 0 0 0 6 10 0 0 52
P2562 vs dcl2
388 303 64 240 275 0 0 0 0 0 11 5 0 0 80
P3611 vs dcl2
297 206 51 228 270 0 0 0 0 0 6 2 0 0 49
P3612 vs dcl2
229 223 53 303 321 0 0 0 0 0 2 2 0 0 53
P4041 vs dcl2
98 155 44 294 299 0 0 0 0 0 2 2 0 0 44
P4042 vs dcl2
B
not DE for F4 or dcl2
DE gene for F4 or dcl2
DE gene for F4 and dcl2
r=.87
p=2.30e-27
rnaseq_dcl2_logFC
rnaseq_P1512_logFC
r=.82
p=1.15e-5
rnaseq_P2561_logFC
r=.68
p=1.19e-16
rnaseq_P2562_logFC
r=.74
p=2.76e-29
rnaseq_P3611_logFC
r=.76
p=2.02e-21
rnaseq_P3612_logFC
r=.83
p=1.98e-29
rnaseq_P4041_logFC

**Extended data Figure 7. Differentially expressed genes are shared between the F4s and *dcl2* mutant.** **a.** Venn diagram of genes upregulated (+DEG) and downregulated (-DEG) for an F4 plant compared to the genes upregulated (+D2G) or downregulated (-D2G) in the *dcl2* mutant. **b.** Scatter plot of the gene expression log fold change (FC) for each F4 compared to the *dcl2* mutant. Grey: non-DEG or in between for any of the two plants; green: DEG for the F4 or *dcl2*; purple: DEG for F4 and *dcl2*. Pearson Correlation coefficient values (r) for the DEG and D2G genes (purple genes) for each scatter plot, p = p value.
